# Does ecology shape geographical parthenogenesis? Evidence from the facultatively parthenogenetic stick insect *Megacrania batesii*

**DOI:** 10.1101/2024.04.02.587815

**Authors:** Soleille Miller, Daniela Wilner, Jigmidmaa Boldbataar, Russell Bonduriansky

**Affiliations:** Evolution & Ecology Research Centre, School of Biological, Earth and Environmental Sciences, UNSW Sydney, Sydney, New South Wales, Australia

**Keywords:** Geographic parthenogenesis, plant-insect interactions, host plant, asexual reproduction, frozen niche variation, general purpose genotype, Benstonea, Pandanus, Megacrania

## Abstract

Closely related sexual and parthenogenetic species often show distinct distribution patterns, known as geographical parthenogenesis. These patterns, characterized by a mosaic of separate sexual and parthenogenetic populations across their natural range, can also be found in facultative parthenogens – species in which every female is capable of both sexual and parthenogenetic reproduction. The underlying mechanisms driving this phenomenon in nature remain unclear. Features of the habitat, such as differences in host plant phenotypes or niche breadth, could favour sexual or asexual reproductive modes and thus help to explain geographical parthenogenesis in natural insect populations. *Megacrania batesii* is a facultatively parthenogenetic stick insect that displays geographical parthenogenesis in the wild. We aimed to explore whether sexual and parthenogenetic populations of *M. batesii* displayed niche differentiation or variations in niche breadth that could explain the separation of the two population types. To do this, we sampled host plants from across the range of *M. batesii* and quantified phenotypic traits that might affect palatability or accessibility for *M. batesii*, including leaf thickness, toughness, spike size and density, height, and chemical composition. We also quantified host plant density, which could affect *M. batesii* dispersal. We found little evidence of phenotypic differences between host plants supporting sexual versus asexual *M. batesii* populations, and no difference in host-plant density or niche breadth between the two population types. Our results suggest that habitat parameters do not play a substantial role in shaping patterns of geographical parthenogenesis in wild populations of *M. batesii*. Instead, population sex ratio variation could result from interactions between the sexes or dispersal dynamics.

## Introduction

In the wild, parthenogenetic and closely related sexual species often show a pattern of differing reproductive modes over a geographic landscape, known as geographic parthenogenesis (Vandel 1928; Lynch 1984). Parthenogenetic lineages are often found in habitats at the edge of the range (Bell 1982) or at higher latitudes or altitudes (Verduijn et al. 2004) and have been associated with habitats affected by Pleistocene glacial cycles (Glesener and Tilman 1978; Suomalainen et al. 1987; Hörandl 2009). However, such patterns do not occur in all parthenogenetic organisms, and their interpretation varies across studies (reviewed in Tilquin and Kokko 2016). Thus, there is no accepted general explanation for why these patterns might arise and be maintained in natural populations.

Differences in ecological niches available to sexual and parthenogenetic animals could explain the geographic distribution patterns of reproductive modes seen in the wild, helping to shed light on the evolution of reproductive modes (Case and Taper 1986; Halkett et al. 2006; Lehto and Haag 2010; Meirmans et al. 2012). Facultative parthenogens - organisms that can reproduce both sexually and parthenogenetically – offer a convenient system to better understand geographical parthenogenesis by investigating ecological niches available to sexually versus parthenogenetically produced conspecifics. While the literature on geographic parthenogenesis has mostly focused on separate obligate sexual and parthenogenetic species (reviews on GP: Kearney 2005; Hörandl 2009), there’s evidence that some facultative parthenogens exhibit similar geographic patterns with distinct sexual and asexual populations across their respective ranges (Morgan-Richards et al. 2010; Burns et al. 2018; Miller et al. 2024). Exploring the environmental factors shaping geographic parthenogenesis in a facultatively parthenogenetic species could enhance our understanding of how ecological factors might influence the evolution of reproductive modes in nature while removing factors that may be due to speciation.

Several theories have been developed to explain the reproductive mode variation found in natural populations of parthenogenic and related sexual animals (reviewed in Vrijenhoek and Parker 2009; Tilquin and Kokko 2016). In particular, it has been hypothesized that, in comparison to sexual lineages, parthenogens could evolve generalist strategies that could enable them to colonize marginal, extreme, or changing habitats (i.e. a “general purpose genotype”; Lynch 1984). Alternatively, it has been hypothesized that parthenogens may experience selective sweeps that create populations of clones that are highly specialized for specific environments (i.e. a “frozen-niche” mechanism; Vrijenhoek 1979). Attempts to test these alternative predictions have mainly focused on study systems that are apomictic, hybrids (which are often also polyploid), or automictic parthenogens that reproduce via central fusion (Bierzychudek 1989; Oplaat and Verhoeven 2015; Van Der Kooi et al. 2019; Hersh 2020). These reproductive pathways often lead to stable or even increased heterozygosity when compared to sexual relatives (Suomalainen et al. 1987; Marescalchi and Scali 2003; Jaron et al. 2021). By contrast, few studies have investigated automictic parthenogens with diploidy restoration mechanisms that result in very low heterozygosity and allelic diversity (e.g. parthenogenesis via terminal fusion or gamete duplication). It therefore remains unclear how automictic parthenogens compare ecologically to their sexual counterparts that have higher genetic diversity (see Larose et al. 2018).

The Peppermint Stick Insect, *Megacrania batesii*, is a facultative parthenogen with a restricted Australian range in tropical north Queensland rainforest (Cermak and Hasenpusch 2000). Every female is capable of both sexual and parthenogenetic reproduction, but with slightly reduced hatching success when reproducing parthenogenetically (Wilner et. al., in prep.). Parthenogenetic reproduction in this species occurs via automixis, and results in very low or zero heterozygosity in offspring (Miller et al., in prep.). In the wild, despite the capacity of each female to switch between reproductive modes, there are distinct mixed-sex and all-female populations of *M. batesii* littered across the range (Miller et al. 2024). Mixed-sex populations typically exhibit an equal sex ratio, and the individuals there mainly undergo sexual reproduction; all-female populations consist exclusively of females, reproducing parthenogenetically. The distribution patterns of these populations resemble patterns of geographic or peripheral parthenogenesis in other species: all-female *M. batesii* populations are often found at the edges of the range, while mixed-sex populations are often found in the center (Miller et al. 2024). *Megacrania batesii* offers an opportunity to test predictions about the ecology of natural populations of parthenogens with extremely low levels of genetic diversity, and to directly compare such populations to sexual populations of the same species.

In their natural range in Australia, *M. batesii* can be found on several species of host plant belonging to two genera, *Pandanus* and *Benstonea* (Miller et al 2024). In addition to food, the host plants also offer protection to *M. batesii*, which rest within the median grooves of the leaves during the day (see Fig. 1E), where they are both camouflaged and shielded from predators by spikes on the edge and bottom midline of each leaf. Both *Pandanus* and *Benstonea* are monocots within the *Pandanacae* family, and there are at least 5 species of *Pandanus* and at least 2 species of *Benstonea* within *M. batesii*’s range (Atlas of Living Australia 2023). However, given high phenotypic variation within each host-plant species and the lack of a full molecular phylogeny for this group, as well as a history of hybridization, it is difficult to identify these plants to the species level (see Buerki et al. 2012). The known range of *M. batesii* in Australia spans ∼250km of Queensland coastline, with most known populations clustered within an area spanning ∼35km of coastal rainforest. Habitats across the range of *M. batesii* show little variation in temperature, precipitation, or altitude, but we noticed that the host plants show considerable morphological variation. We also noticed much variation in the density and distribution of host plants between habitats occupied by different *M. batesii* populations. Some *M. batesii* populations are found in high-density habitats where host plants are smaller (immature) individuals that occur in contiguous patches, often with overlapping leaves. Other populations are found in low-density habitats where many of the host plants are trees located several meters apart. Denser habitats may facilitate dispersal in this species as *M. batesii* are flightless and can only disperse by walking short distances between host plants (Boldbaatar 2022). Low-density habitats may therefore favour the establishment of all-female populations because only one female would need to reach the new host plant to colonize that area; whereas both a male and female would need to reach a habitat patch to form a new mixed-sex population. Thus, small-scale environmental heterogeneity in the form of host plant phenotypic and distributional variation could contribute to the pattern of geographic parthenogenesis observed in natural populations of *M. batesii*.

**Figure 1:**
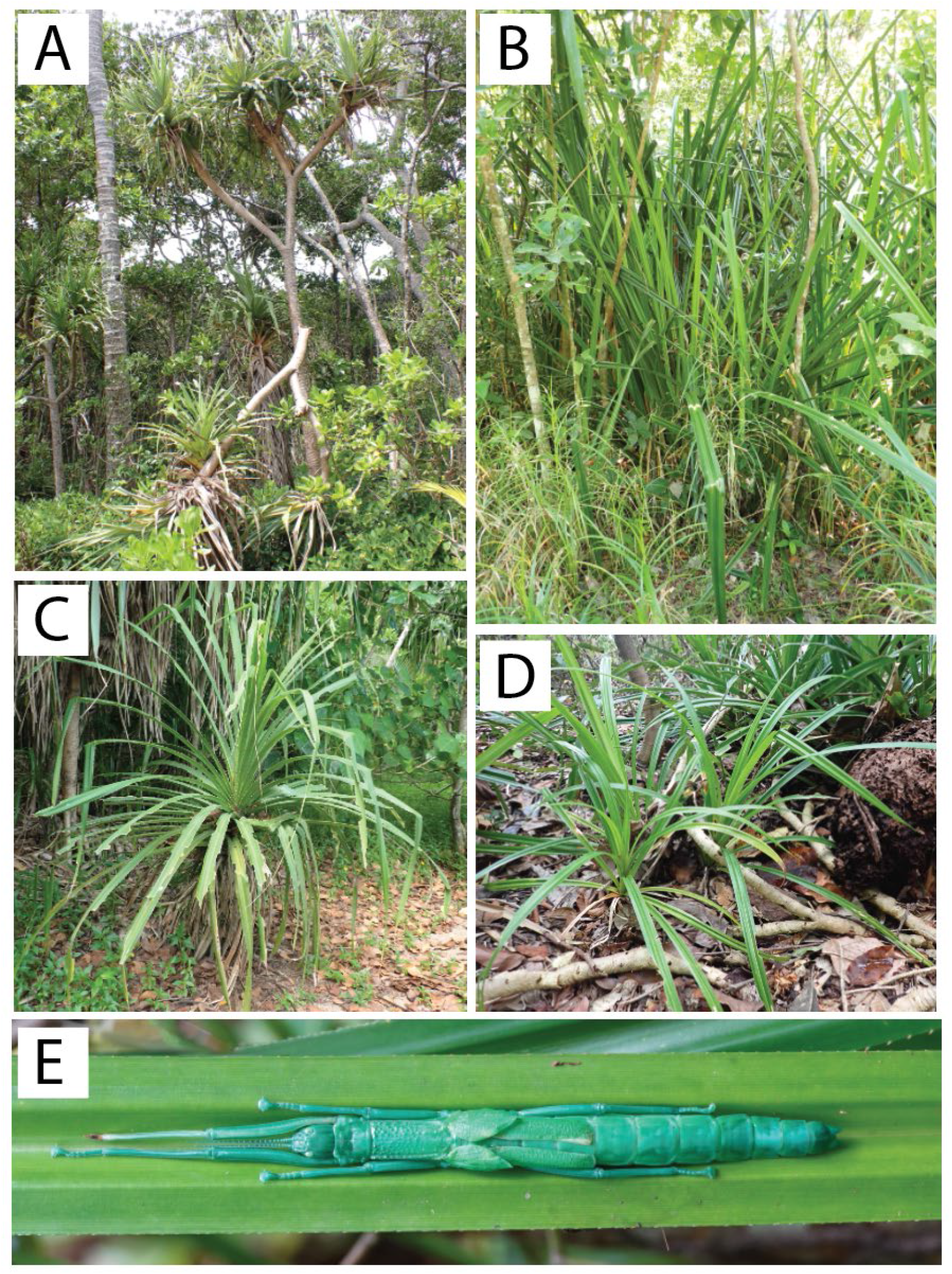
Host plants used by *Megacrania batesii*: A) a mature *Pandanus tectorius* plant with a woody trunk, B) a *Benstonea lauterbachii* plant growing in a swamp C) an immature *Pandanus sp.* plant, D) *Benstonea monticola* plants, and E) *M. batesii* female resting in the groove of a *Benstonea monticola* leaf.

We also observed that certain host plants appear to remain completely untouched by *M. batesii*, despite being located adjacent to, and sometimes even in contact with, similar-looking plants that have been heavily consumed by *M. batesii*. This suggests that there could be morphological or chemical differences between eaten and uneaten host plants. For example, plants may remain uneaten if they have thicker leaves or larger spikes that curb herbivory. Leaf thickness and toughness are common defences against herbivores and are often negatively correlated with herbivory (Peeters 2002; Kursar and Coley 2003; Fine et al. 2004; Clissold et al. 2009; Pringle et al. 2011; Caldwell et al. 2015). *Pandanus* and *Benstonea* species used as host plants by *M. batesii* possess spikes that line the edge of the leaves, and these spikes may also function as a deterrent against herbivory by *M. batesii* as they initiate feeding at the edge of the leaf (Cermak and Hasenpusch 2000). Specifically, the length of the spikes, the angle at which they protrude, and the density of the spikes may play a role in protecting the host plants against attacks by *M. batesii*. Some of the host plants we observed in the field were tall, mature trees, with edible leaves only occurring at the top of the tree (see Fig. 1A) whereas other host plants were immature or growing as shrubs lacking a woody trunk, providing more easily accessible leaves (see Fig. 1C and 1D). Thus, host-plant height could also affect *M. batesii* herbivory as taller trees may impede *M. batesii* dispersal to the section of the tree with edible leaves, requiring the insects to be exposed to predation for longer to get to their food source. Additionally, many plants produce chemical defences against herbivorous insects, with high carbon to nitrogen ratios (C/N) being associated with accumulation of carbon-based phenolics which can reduce herbivory (Bryant et al. 1983; Lindroth et al. 2001; Wittstock and Gershenzon 2002; Fürstenberg-Hägg et al. 2013). Nitrogen content is especially important in plant-insect interactions as protein and nitrogen concentrations are positively correlated in leaves (Throop and Lerdau 2004), and studies have shown that nitrogen concentrations directly affect growth and diet choice in many herbivorous insects, leading to insect preferences for higher nitrogen in leaves (Mattson 1980; Rausher 1981; Osier and Lindroth 2001). Thus, host plants may vary in morphological or chemical traits that affect palatability for *M. batesii*.

In this study, we aimed to quantify and compare host-plant phenotypes from habitats hosting sexual versus parthenogenetic stick insects in their natural range. Specifically, we aimed to understand whether parthenogenetic *M. batesii* show patterns consistent with the general purpose genotype theory or the frozen niche variation theory to explain the geographic parthenogenesis observed in the wild. The general purpose genotype theory predicts that parthenogenetic stick insects inhabit habitats with host plants that are more morphologically or chemically diverse. This would suggest that parthenogenetic *M. batesii* lineages have a wider niche breadth than their sexual counterparts, reflecting broader tolerance for morphological and chemical variation in their host plants. Alternatively, the frozen niche theory predicts that parthenogens are highly specialised with a narrow niche breadth, and therefore all-female populations of *M. batesii* will be found in habitats with little diversity of phenotypic and chemical traits, possibly with different but similarly narrow niches for each all-female population. Additionally, we aimed to test the prediction that asexual populations are more likely to establish and persist in low-density habitats. Lastly, we asked whether there were chemical or morphological differences between host plants that were eaten versus host plants that were avoided entirely by *M. batesii*, and whether parthenogenetic *M. batesii* have greater or lesser tolerances for host plant morphological and chemical defences. To address these questions, we quantified variation in host plant leaf morphology (leaf thickness, toughness, and slenderness, spine length and orientation,) and chemistry (carbon, hydrogen and nitrogen content), and host plant height and density (in 10x10m quadrats) in several mixed-sex and all-female populations of *M. batesii* within their natural range in far-north Queensland.

## Methods

We studied the habitats of mixed-sex (sexual) and all-female (parthenogenetic) populations of *M. batesii* along ∼35 km of coastline between Cape Kimberley and Cape Tribulation, Queensland (see Fig. 2). We also investigated two isolated all-female populations located ∼ 160-180 km south of the Daintree River at Etty Bay and Bingil Bay. These *M. batesii* populations are described in Miller et al (2024), except for one population (MAD). Habitat for *M. batesii* was identified as aggregations of host plants of the genus *Pandanus* or *Benstonea* (or both). Mixed-sex populations of *M. batesii* were identified as having an approximately equal sex ratio of males to females, and sex ratios of hatchlings from eggs collected from such populations were approximately even, confirming that reproduction is predominantly sexual. Conversely, all-female populations were identified as having only females, and hatchlings from eggs collected from such populations were all females, confirming that reproduction is exclusively parthenogenetic. One “transitional” population appears to be in the process of transitioning from an all-female population of *M. batesii* to a mixed-sex population (see Miller et. al., 2024). Host plants from the transitional population were excluded from analyses comparing all-female and mixed-sex populations as this population cannot be included in either of those two groups. However, this population was included in the analyses comparing eaten and uneaten host plants because *M. batesii* population type was not relevant to that question. We collected phenotypic data on a total of 89 *Pandanus* sp. or *Benstonea* sp. plants, including 27 plants from 6 mixed-sex populations, 49 plants from 8 all-female populations, and 13 plants from the transitional population. Chemical data was collected from 43 of these plants. We also collected density data on a total of 28 quadrats, 14 from sites where mixed-sex populations occur, and 14 from sites where all-female populations occur (see Fig. 2 and Table 1).

**Figure 2:**
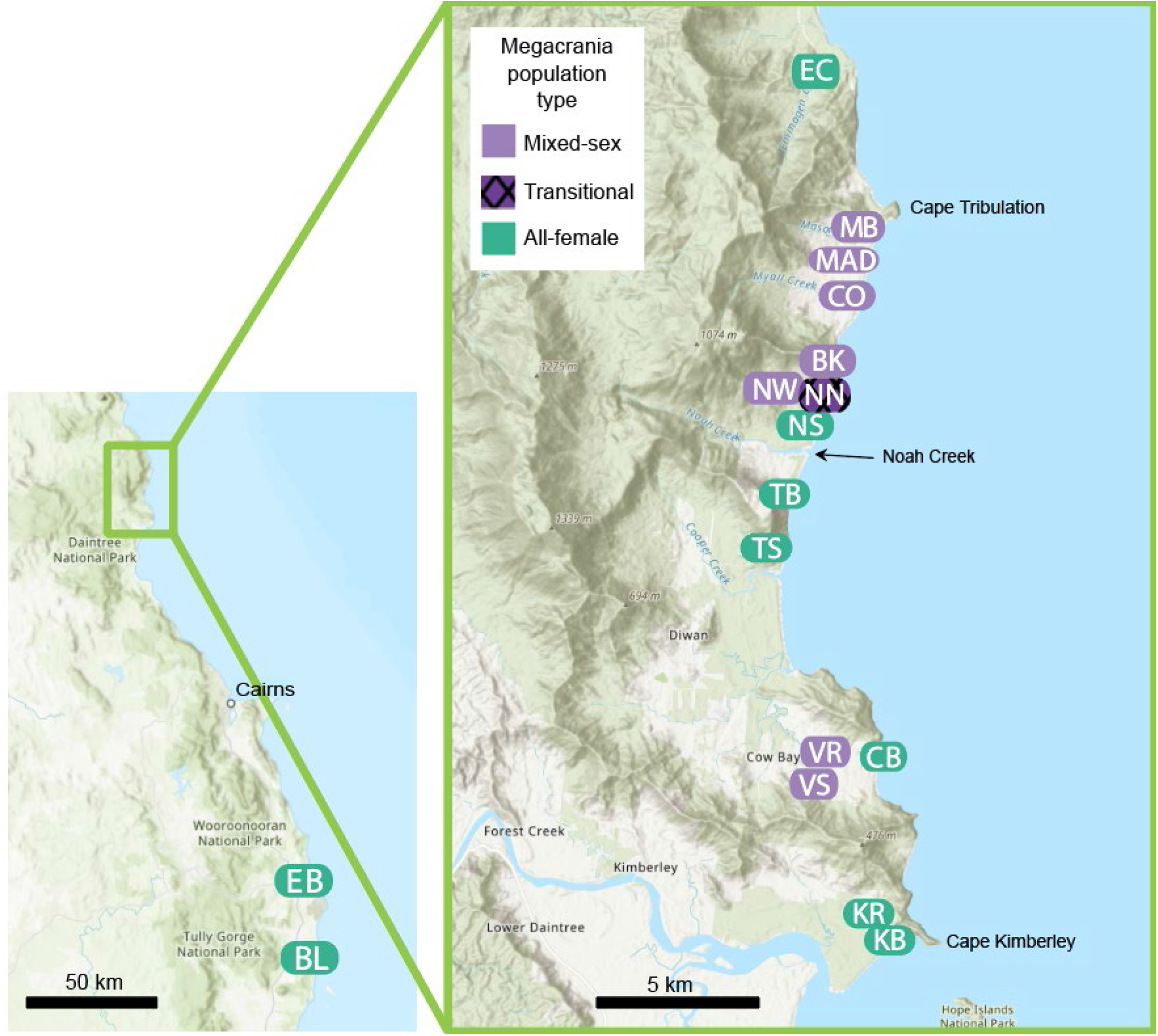
Habitat sampling sites sampled in this study. The sampled sites are a subset of the sites containing *Megacrania batesii* populations studied by Miller et. al. (2024) with the addition of one new population (MAD). Color indicates whether the *M. batesii* population at the site is mixed-sex or all-female. The patterned color indicates the transitional population of *M. batesii*.

**Table 1:** Sites and sample sizes for plant and habitat data. Site codes correspond to codes in Miller et al. (2024), apart from MAD. Note: Sample sizes reported here include samples taken from plants with no *Megacrania batesii* chew marks, thus they may differ from sample sizes in analyses where we excluded uneaten plants.

| *M. batesii* site | *M. batesii* population type | No. plants in morphology analyses | No. plants in chemical analyses | No. quadrats for density analyses |
| --- | --- | --- | --- | --- |
| EC | All-female | 7 | 0 | 0 |
| MB | Mixed-sex | 10 | 9 | 4 |
| MAD | Mixed-sex | 5 | 5 | 1 |
| CO | Mixed-sex | 4 | 3 | 3 |
| BK | Mixed-sex | 6 | 0 | 2 |
| NW | Mixed-sex | 3 | 3 | 1 |
| NN | Transitional | 13 | 4 | 0 |
| NS | All-female | 3 | 0 | 2 |
| TB | All-female | 11 | 4 | 3 |
| TS | All-female | 2 | 2 | 1 |
| CB | All-female | 11 | 7 | 2 |
| VR | Mixed-sex | 2 | 2 | 1 |
| VS | Mixed-sex | 2 | 2 | 2 |
| KR | All-female | 2 | 2 | 1 |
| KB | All-female | 0 | 0 | 1 |
| EB | All-female | 3 | 0 | 3 |
| BL | All-female | 5 | 0 | 1 |

To quantify plant morphological traits, we measured plant height using a measuring tape for plants < 2 m in height, and estimated height in m for taller plants. We measured the thickness of three leaves on each sampled plant with a digital micrometer and averaged these three measurements to obtain a leaf thickness value for the plant. We estimated leaf toughness by quantifying the pressure (in kg/cm²) required to cut from the edge to the midline of each leaf using scissors and a penetrometer. To do this, a small divet was drilled into the handle of the scissors. Then, the base of the penetrometer was pressed into the divet until the scissors cut to the midline of the leaf. Three measurements of leaf toughness from the same leaves were obtained in this way for each plant, and the average computed. Thickness and toughness measurements, as well as photos of leaf spikes were taken at ∼30 cm from the base of the leaf when possible, and the oldest and newest leaves on the plant were avoided. We took photos of the leaves of each sampled plant with a scale, and we measured the average length of 5 spikes along the leaf margin from base to tip (mm), as well as the average perpendicular distance between the tip of the spike and the edge of the leaf as a proxy for the angle of the spike (mm), and average distance between spikes (mm) using ImageJ (Schneider et al. 2012). Photos of the entirety of each plant were used to calculate the average leaf slenderness ratio (leaf length/leaf width) on ImageJ, measuring three representative leaves per plant.

To quantify plant chemical traits, we obtained leaf clippings (∼3 x 2 cm) from 39 of the sampled plants using scissors. The blades of the scissors were sterilised with ethanol prior to taking each sample. 19 clippings were obtained from habitats hosting mixed-sex populations of *M. batesii* and 20 clippings were obtained from habitats hosting all-female populations; no clippings were taken from the transitional *M. batesii* population. The clippings were dried at 55°C for 48 hours using a Thermoline Scientific Laboratory Oven (model: TEO-66F; https://www.thermoline.com.au/) at the JCU Daintree Rainforest Observatory (https://www.jcu.edu.au/daintree). Dried leaf samples were then sent to the Mark Wainwright Analytical Center at UNSW (https://www.analytical.unsw.edu.au) for analyses of carbon, hydrogen, and nitrogen concentrations (i.e., percentage composition) in each sample.

To quantify density of host plants in sample sites across the range, we sampled 10 x 10 m quadrats from 7 sites hosting mixed-sex populations of *M. batesii* and 9 sites hosting all-female populations of *M. batesii*, with some sites having multiple quadrats (see table 1). Quadrats were sampled only in habitat patches with conspicuous herbivory by *M. batesii*. The chew-marks of *M. batesii* on *Pandanus* and *Benstonea* host plants are readily distinguishable from leaf damage caused by other herbivores: *M. batesii* feeds by making distinctive, elongated cuts along the leaf edge (see Cermak and Hasenpusch 2000). No quadrats were taken from habitat hosting the transitional *M. batesii* population. Within each quadrat, we counted the number of host plants (*Pandanus* and *Benstonea*) and divided by 100 to estimate the number of plants per square meter. Most of the quadrats sampled consisted largely of either *Pandanus* sp. or *Benstonea* sp. plants. Other vegetation within the quadrats typically consisted of grasses or shrubs that are not used by *M. batesii*.

All analyses and data plots were done using R Statistical Software (v4.2.2; R Core Team 2022). Morphological analyses were based on measurements of plant height, average leaf thickness, average leaf toughness, spike angle (log-transformed to meet the assumption of normality), average distance between spikes, average spike length, and leaf slenderness ratio. Chemical analyses were based on nitrogen, hydrogen, and carbon content of leaves, expressed as percentages of sample mass. For both chemical and morphological analyses, we ran Principal Components Analyses (PCA) using data that was centred, scaled and then analysed using the function *prcomp()* from the package “vegan” (Dixon 2003). Trait loadings and individual scores on the first two principal components were then visualized using the function *ggbiplot()* from the package “ggbiplot” (https://github.com/vqv/ggbiplot).

Both morphological and chemical data was first standardized to account for variation in scale between the different variables. This was done using the *vegan* function *decostand()* with the argument *method=”standardize”*. This created a new multivariate object with a mean of zero and variance of one for each variable. Then, a Euclidean distance matrix was created using the function *vegdist().* We ran a permutation regression model (999 permutations) using this matrix as the response variable and the presence/absence of *M. batesii* chew marks as well as host-plant genus (to account for genera differences between host plants) and their interaction as fixed predictor variables with the function *adonis2()*. This permutation model partitions sums of squares of a multivariate data set and is analogous to multivariate analysis of variance (Anderson 2001). To investigate the interaction between presence/absence of *M. batesii* chew marks and host-plant genus, we first separated genera and ran the PCA visualizations again for each genus. We paired this with two more permutational models (one for each genus) in *adonis()* to extract the coefficients and understand how each plant trait was affecting palatability in each genus. To understand how C/N ratios affected herbivory by *M. batesii*, we used a linear model with C/N ratio as the response variable and presence/absence of *M. batesii* chew marks as well as host-plant genus (and their interaction) as fixed predictor variables.

Density of host plants in each quadrat (plants/m^2^) was log-transformed to meet the assumption of normality. A linear mixed model regression was performed on the transformed data using the function *lmer()* from the lme4 package (Bates et al. 2015), with density as the response variable, *M. batesii* population type (mixed-sex or all-female) as the fixed effect, and site as a random effect.

Data collected from the transitional *M. batesii* habitat, as well as plants that were not found to have *M. batesii* chew marks, were excluded from analysis of niche differences, leaving 62 samples for this analysis. To understand whether *M. batesii* from all-female populations (parthenogenetic) occupy different niches in terms of morphological and chemical host plant traits than their sexual counterparts, we conducted a PCA on the same set of morphological and chemical traits as our analysis on *M. batesii* preferences. Again, this data was centred, scaled, and then analysed with a PCA, using the function *prcomp()*. Trait loadings and individual scores on the first two PCs of both PCAs were then visualized using the function *ggbiplot().* To test whether there were differences in host plant chemical and morphological traits between host plants in mixed-sex vs all-female *M. batesii* populations, we ran the same permutation regression model as described above for *M. batesii* preferences. However, we adjusted the predictor variable from presence/absence of *M. batesii* chewmarks to *M. batesii* population type (mixed-sex or all-female).

We then tested for niche breadth differences, specifically whether a particular reproductive mode was found at sites where host plants exhibited a wider range of morphological or chemical traits. We first wanted to see if the host plants harbouring all-female *M. batesii* populations exhibit more consistent phenotypes across their range compared to host plants harbouring mixed-sex *M. batesii* populations. This would show us whether asexual *M. batesii* are more likely to persist in habitats with a specific host plant phenotype. To do this, we first used the *betadisper()* function, which calculates distances between group members (host plant phenotypes) and centroid of the group (whether that plant was hosting a mixed-sex or all-female population of *M. batesii*) by reducing the original Euclidean distances to principal coordinates, an analogue of Levine’s test for homogeny of variants, but in a multivariate space (O’Neill and Mathews 2000; Anderson 2001). We did this for morphological and chemical traits separately. Then, to see if all-female populations of *M. batesii* had a wider or narrower niche overall (i.e., across sites), we ran a permutational analysis using the function permutest() with default settings. This function compared the average group dispersions between phenotypes of host plants found in all-female versus mixed-sex *M. batesii* populations using a *t*-test, and then performed a permutation test based on the *t* statistic derived from that pairwise comparison. We also did this for the morphological and chemical traits separately.

Next, we to see if habitats hosting all-female *M. batesii* populations had lower within-site host plant phenotypic diversity (i.e. possibly different, but similarly narrow niches) than habitats hosting mixed-sex *M. batesii* populations. Similarly to the last analysis, we ran *betadisper()* again, this time with the group being site (e.g. CB, BK, TB, etc.) for both the chemical and morphological data sets. We modelled the distances to the within-site centroid using a linear mixed effect model with *M. batesii* population type as a fixed effect, and site as a random effect. The distance to the group centroid (i.e. habitat site) was log-transformed to meet the assumption of normality. We did this for the morphological and chemical traits separately using the *lmer()* function. All visualizations were made using *ggplot2*.

## Results

In sites supporting *M. batesii* populations, *Pandanus* and *Benstonea* plants were found in two distinct environments. *Pandanus* spp. plants were typically found growing in coastal rainforest margins, whereas *Benstonea* spp. plants typically grew in and around streams and swamps in closed-canopy rainforest. Habitats hosting *M. batesii* typically included either *Pandanus* spp. or *Benstonea* spp. plants, but rarely both genera in close proximity. Generally, *Pandanus* plants were taller and spread further apart, whereas *Benstonea* plants were relatively smaller and spaced closer together (sometimes overlapping). However, we found much variation between sampled sites in terms of host-plant morphology and density. For example, at some sites we found patches of many small *Pandanus* plants growing close together; while at other sites we found mature *Pandanus* trees growing meters apart. In some swamp sites, we found many small- and medium-sized *Benstonea* plants (with and without obvious woody trunks) growing close to each other (with overlapping leaves) in muddy substrate near but not immersed in water; while in others we found large *Benstonea* plants growing in stream or swamp water, either in clumps of multiple plants or as isolated individuals.

Plants of the family *Pandanaceae* are notoriously difficult to classify to species level, due to several key morphological traits having evolved independently and a rich history of hybridization (Buerki et al. 2012). Consequently, we were unable to confidently determine species identity of each plant based on morphology and did not include a species-level phylogeny in our analysis. However, based on records in the Atlas of Living Australia, there are at least 5 species of *Pandanus* in the sampled area (*Pandanus tectorius*; *Pandanus spiralis*; *Pandanus solms-laubachii*; *Pandanus gemmifer*; *Pandanus cookii*), the most common being *Pandanus tectorius.* There are also at least 2 species of *Benstonea* (*Benstonea monticola*; *Benstonea lauterbachii*), the most common being *Benstonea monticola*. These two *Benstonea* species have clearly distinct morphologies when mature (*B. lauterbachii* having much longer leaves and tending to grow directly in water) but are difficult to distinguish when immature. The genera *Pandanus* and *Benstonea* are clearly distinct morphologically, and we include genus identity for all sampled plants in the analyses reported below. *Pandanus* plants have relatively thicker and shorter leaves, with larger spikes, and grow as trees > 3 m high when mature. *Benstonea* plants have relatively thinner, longer leaves with smaller spikes, and can develop thin woody trunks ∼1-3 m high (see Fig. 1; Callmander et al. 2012).

*Benstonea lauterbachii* (Fig. 1B) was found in multiple sites where *M. batesii* occurred, but mature *B. lauterbachii* plants never showed signs of being consumed by *M. batesii*. Other species, especially *Benstonea monticola* and *Pandanus tectorius*, were frequently used by *M. batesii* as host plants. However, in these and other host-species, certain individual plants were completely untouched by *M. batesii* despite being located adjacent to, and sometimes even in contact with, morphologically similar and presumably conspecific plants that had been heavily consumed by *M. batesii*. Based on the set of morphological traits measured (plant height, leaf thickness, leaf toughness, spike length, spike angle, distance between spikes, slenderness ratio), there were no clear differences in morphology between eaten and uneaten plants (See Fig. A1), with PCA showing nearly complete overlap between the two groups (See Fig. 3A). Statistical analyses showed a significant interaction between genus and palatability (i.e., whether the plant showed evidence of feeding by M. batesii; PERMANOVA; p=0.034; Table A1a). Further investigation into this interaction showed five of the seven morphological traits that we quantified had effects of opposite sign on palatability of Pandanus versus Benstonea plants (See Fig. A2; Table A2). This suggests that the plant traits that affect palatability for *M. batesii* differ between *Pandanus* vs *Benstonea spp*., or they exhibit varying degrees of importance depending on genus. By contrast, we found clear differences in chemical composition between host plants that showed evidence of *M. batesii* feeding versus host plants that were untouched by *M. batesii* (PERMANOVA; p<0.001; See Fig. 3B; Table A1b). Eaten host plants had higher levels of carbon and hydrogen and lower levels of nitrogen than uneaten plants (see Fig. 4). Likewise, eaten plants had higher C/N ratios when compared to uneaten plants (Table A1c and Fig. 4D).

**Fig. 3:**
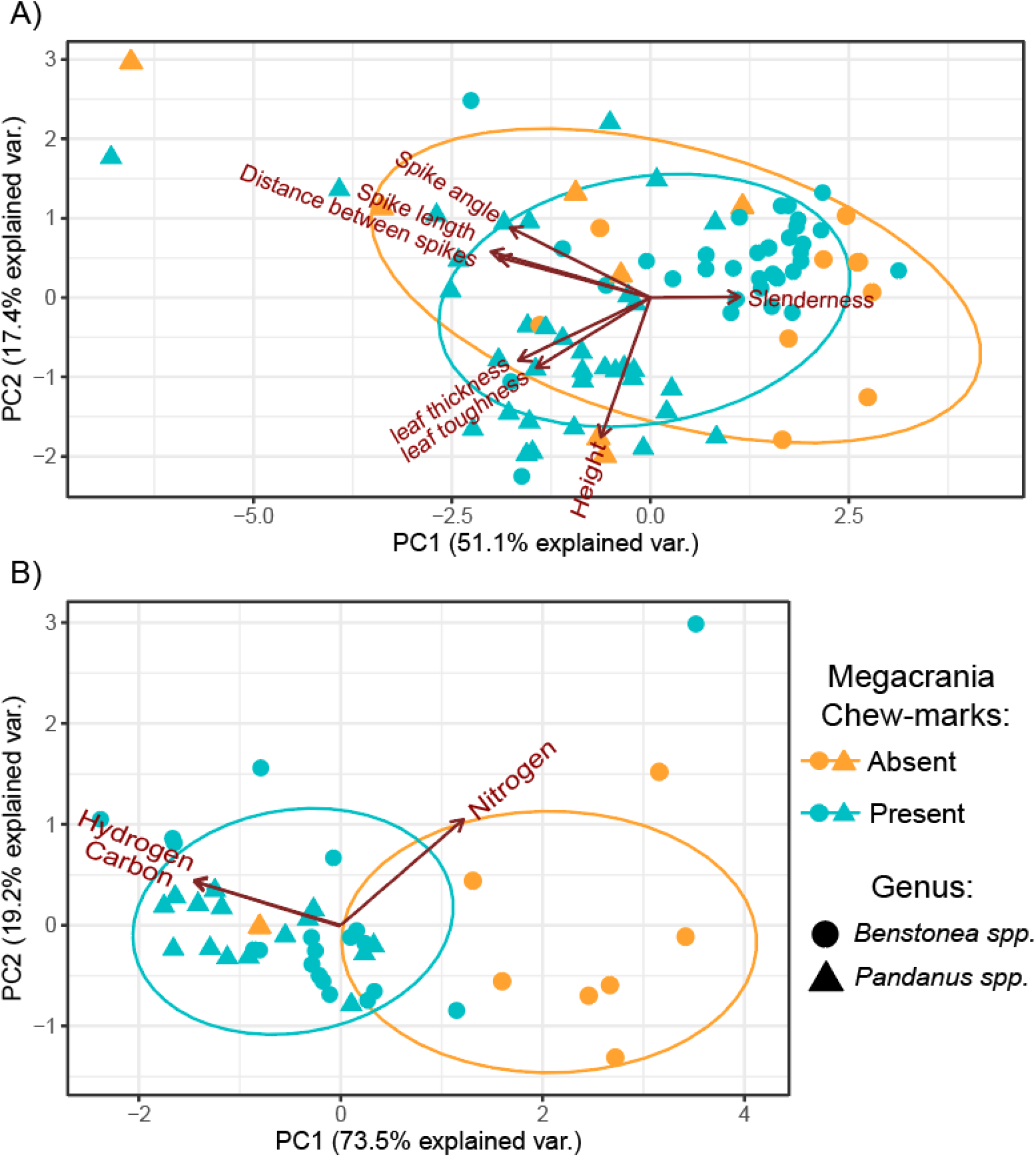
Herbivory on *Pandanus* and *Benstonea spp*. by *Megacrania batesii*. PCA of host plant a) morphological and b) chemical traits. Each point represents an individual host plant sample, the shape indicates whether that sample was a *Pandanus* sp. plant or a *Benstonea* sp. plant, and the color indicates whether *M. batesii* chew-marks were present or absent.

**Figure 4:**
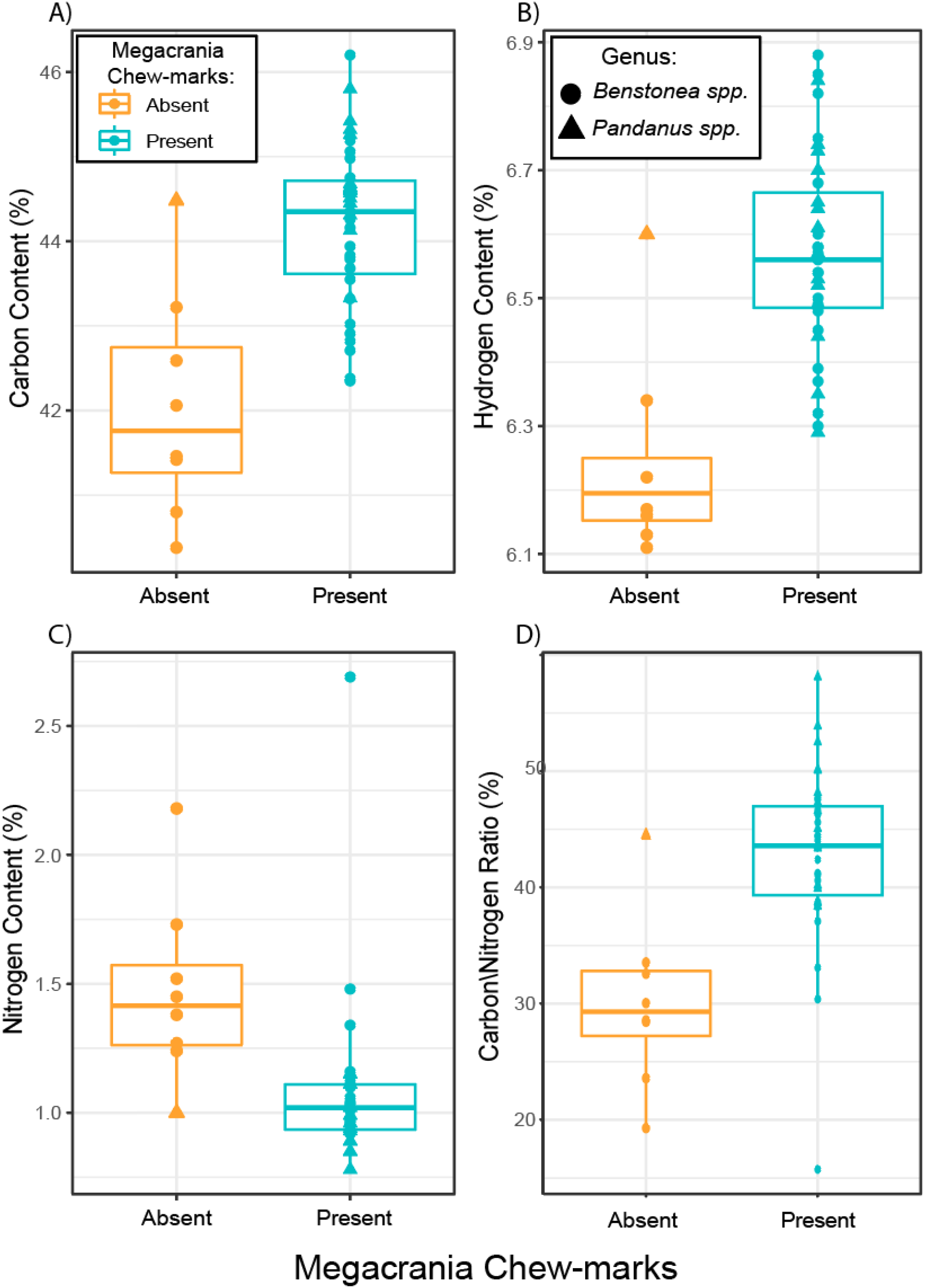
Chemical composition of A) carbon, B) hydrogen, C) nitrogen (%), and D) carbon/nitrogen ratio in individual plants with presence/absence of *Megacrania batesii* herbivory. The lower and upper hinges of the box correspond to the first and third quartiles (25^th^ and 75^th^ percentiles). The median line is shown. The whiskers extend from the hinges to the largest and smallest value within 1.5 times the inter-quartile range. Each point represents a host plant sample, the shape indicates whether that sample was a *Pandanus* sp. or a *Benstonea* sp. plant, and the color indicates whether *M. batesii* chew-marks were present or absent.

*M. batesii* sites varied greatly in density and size of host plants. Some sites were characterized by numerous small- and medium-sized *Pandanus* or *Benstonea* plants densely packed with overlapping leaves, while other sites had large plants (usually, mature *Pandanus* sp. trees) spaced further apart. Density tended to be higher in *Benstonea* habitats (0.53 plants/m^2^) than in *Pandanus* habitats (0.26 plants/m^2^). At one site, many *Benstonea monticola* plants grew ∼50 cm apart, with heavily overlapping leaves. We found no evidence for a difference in host-plant density between sites occupied by all-female versus mixed-sex *M. batesii* populations (see Fig. 5; linear mixed model; p=0.661; Table A3).

**Figure 5:**
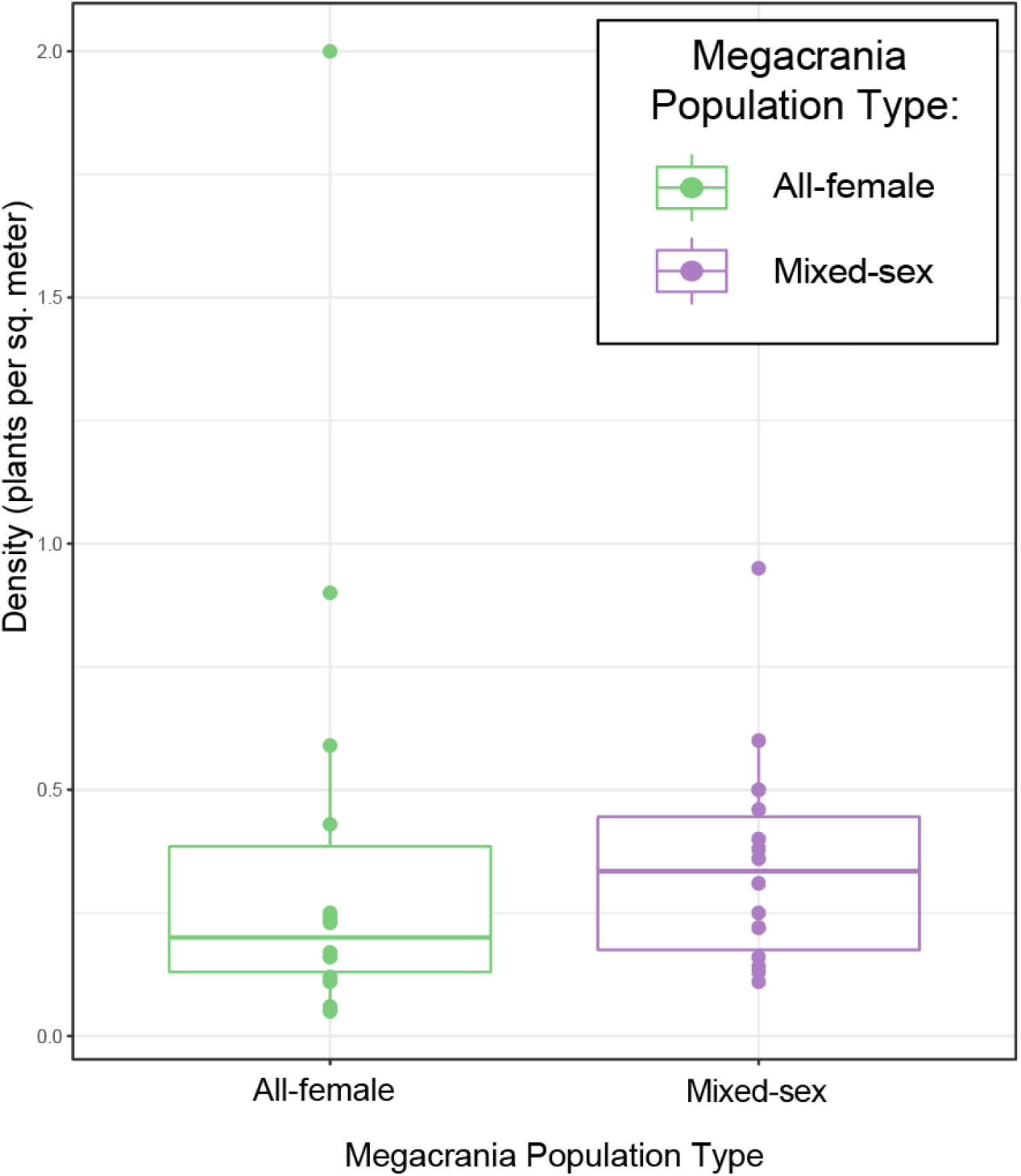
Host plant density in 10 x 10 m quadrats sampled in habitats occupied by all-female or mixed-sex *Megacrania batesii* populations. Each point is one quadrat sampled and the color indicates whether that quadrat hosted an all-female population or a mixed-sex population of *M. batesii*. The lower and upper hinges of the box correspond to the first and third quartiles (25^th^ and 75^th^ percentiles). The median line is shown. The whiskers extend from the hinges to the largest and smallest value within 1.5 times the inter-quartile range.

We found some evidence that all-female populations of *M. batesii* used host plants that were morphologically different from the host plants used by mixed-sex populations. PCA showed a pattern of highly overlapping data points on the first two principal components (See Fig. 6A) but permutational modelling revealed a difference between mixed-sex and all-female populations (Table A1d; p=0.015). Relative to mixed-sex populations, all-female populations were more likely to be found on host plants with larger spikes and spikes angled further away from the leaf, as well as thicker and tougher leaves (Fig. A3). Nonetheless, population type explained little variation in plant morphology (R^2^ = 0.04). Comparing chemical traits of plants, we found no differences between plants in habitats occupied by all-female versus mixed-sex populations (see fig. 6B and Table A1e).

**Figure 6:**
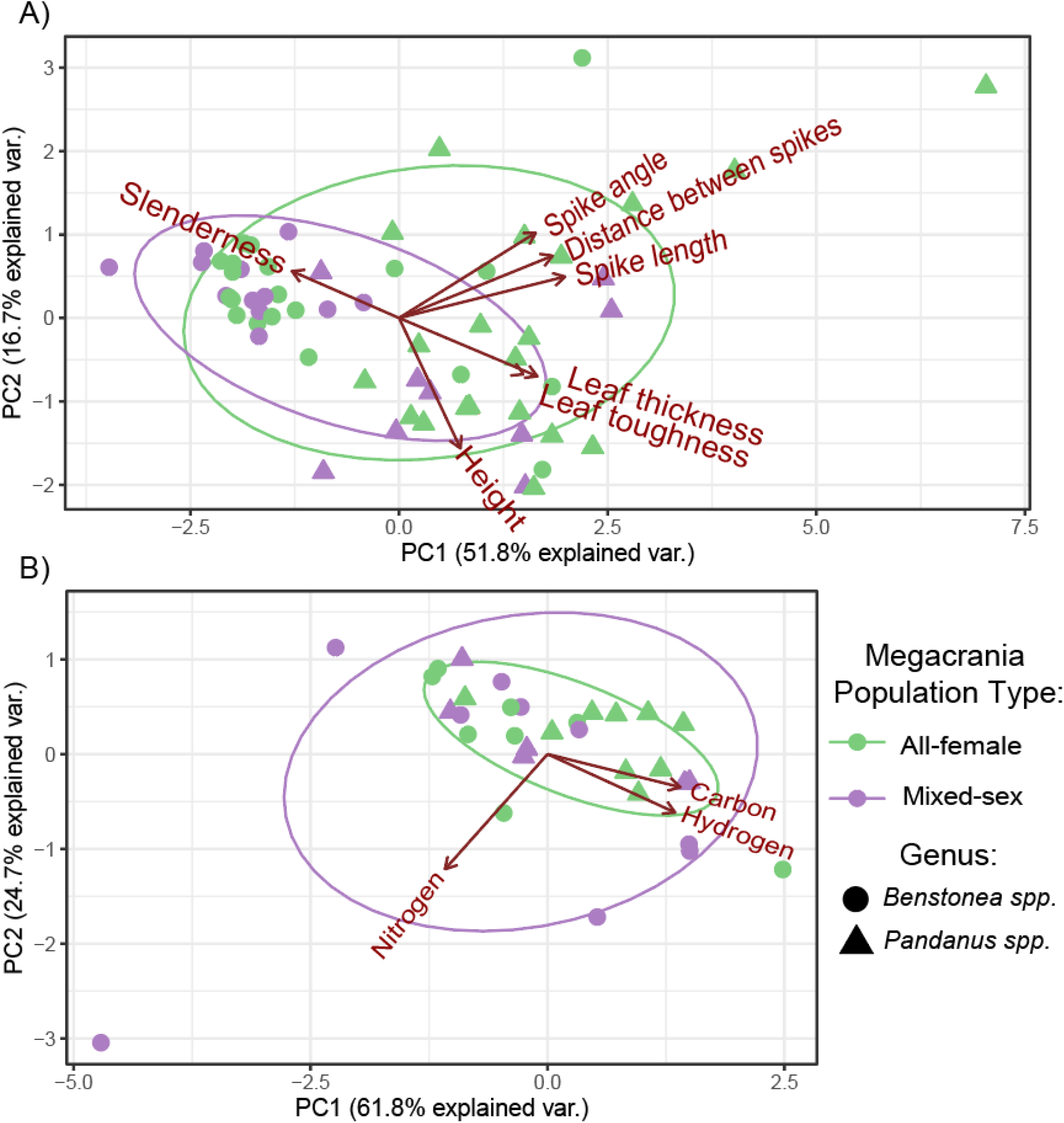
PCA results for host plant a) morphological and b) chemical traits. Each point represents an individual host plant sample, the shape indicates whether the sample is a *Pandanus* or *Benstonea*, and the color indicates whether that plant was within an all-female or mixed-sex *Megacrania batesii* population.

PCA suggested a wider range of morphological trait values of plants hosting all-female populations of *M. batesii*, suggesting that parthenogenetic *M. batesii* occupy a wider niche in terms of host plant morphology across the range (Fig. 6A). Permutational statistical testing, however, did not support this result (Table A4a). The opposite trend was suggested by PCA on chemical traits, where plants hosting mixed-sex populations of *M. batesii* appeared to have a wider range of trait values (see Fig. 6B), but this result was also not supported statistically (Table A4b). Some sites had much within-site variability regarding morphological and chemical traits (see Fig. 7). However, linear mixed-effects modelling did not support a difference in within-site host-plant variability in either morphological or chemical traits between sites hosting all-female vs. mixed-sex *M. batesii* populations (Table A5, Fig. 7). Thus, we found no evidence of an overall difference across all sites in niche breadth between sexual and parthenogenetic *M. batesii* populations.

**Figure 7:**
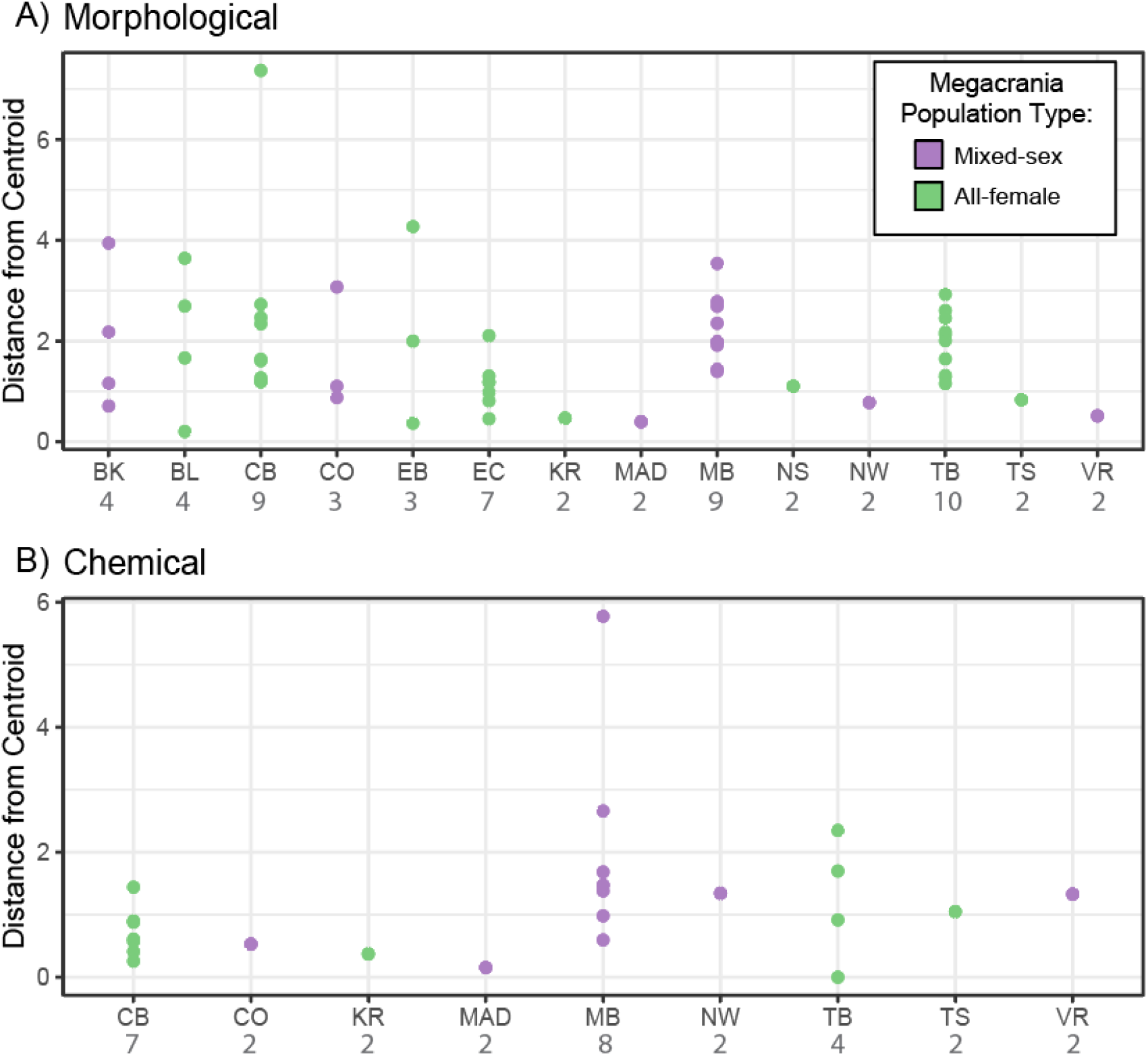
Distance to the group (site) centroid for each sampled plant from each site for a) morphological traits and b) chemical traits. Each point is one host plant and the color indicates whether that site hosts an all-female population or a mixed-sex population of *Megacrania batesii*. Sample sizes for each site are in grey under the site label.

## Discussion

Environmental factors are believed to play a large role in the emergence and maintenance of geographic parthenogenetic patterns in natural populations (Glesener and Tilman 1978; Suomalainen et al. 1987; Tilquin and Kokko 2016), but the factors driving differentiation in reproductive mode among populations of facultatively parthenogenetic animals remain poorly understood. We found some evidence of morphological differences between host plants in habitats inhabited by mixed-sex versus all-female *M. batesii* populations. Specifically, all-female populations utilized host plants with thicker, tougher leaves and larger spikes (see Fig. A3). This could be due to the sexual dimorphism of *M. batesii*: females are substantially larger than males, and might therefore possess a greater capacity to consume host plants with long spikes and thick leaves.

However, while this difference was supported statistically, *M. batesii* population type explained little of the variance in host plant morphology or chemistry. Thus, phenotypic variation among host plants does not appear to exert a substantial influence on the geographical distribution of all-female versus mixed-sex *M. batesii* populations. Host plant density is likely to affect the geographic distribution of *M. batesii* as they rely on their host plants for shelter, safety, and food (Cermak and Hasenpusch 2000). *M. batesii* adults of both sexes are flightless and have been shown to have a very limited capacity for dispersal, so distance between host plants may act as a barrier and a factor influencing the distribution of *M. batesii* populations (Boldbataar 2024). We expected all-female populations to thrive at sites where patches of host plants are more distant from each other (i.e. low-density habitats) because, in such circumstances, the successful arrival of a single female to a distant host plant could initiate the establishment of a new all-female population. Dispersal could occur via walking, rafting on vegetation during a flood event, or dispersal of eggs by ocean currents/flooding. In contrast, the chances of both a male and a female successfully arriving at a new host plant in a low-density habitat patch would be notably lower. Moreover, even if a mixed-sex population arose in a small, isolated habitat patch, stochastic male extinction would be likely to occur eventually, leaving an all-female population (Miller et al. 2024). Our data suggests a weak trend consistent with that prediction (see Fig. 5), but the difference in host plant density between all-female and mixed-sex populations was not supported statistically (Table A3). The quadrat sizes in this study may have been too small to capture the variation between these two groups. It is also possible that the distance between habitat patches is more relevant to this question than distances between host plants within patches. Given the limited mobility of *M. batesii* nymphs and adults (Boldbaatar 2022), the distance between clusters of host plants, which can be tens or hundreds of meters apart, could be a significant barrier to dispersal.

We also investigated whether predictions stemming from the general-purpose genotype (GPG) or frozen-niche variation (FNV) theories could account for the geographic distribution of all-female and mixed-sex *M. batesii* populations. The GPG theory predicts that parthenogenetic animals will have a wider niche breadth whereas the FNV theory predicts a narrower niche breadth in parthenogenetic animals compared to their sexual counterparts. We tested niche breadth generally in plants that supported parthenogenetic versus sexual *M. batesii* populations (overall host-plant variability). We also tested niche breadth within each *M. batesii* population site (within-site host-plant variability) and compared the all-female to the mixed-sex population sites. If each all-female site contains a different specialist genotype, overall host-plant variability could be high for parthenogenetic *M. batesii*, but within-site variability would be low by comparison with sexually reproducing *M. batesii*. We found little evidence of differentiation in either overall or within-site host-plant variability in morphological or chemical traits (see Table A4 and A5), suggesting that niche breadth does not differ substantially between parthenogenetic and sexual *M. batesii* populations. It is possible that the distributions of AF vs MS populations could be explained by other environmental factors that form part of the niche, such as temperature, precipitation, or humidity. However, *M. batesii* inhabits a tropical environment characterized by relatively uniform climactic conditions across its limited central range (latitude difference = 0.23°), and both population types occur throughout the central part of the range of *M. batesii* in Australia (although BL and EB, the two isolated populations at the southern extreme of the range, ∼1.25° south of the other populations, are both all-female). Large-scale climate parameters are therefore unlikely to play a major role in the distribution of mixed-sex versus all-female populations; although there could be unmeasured microclimate differences between sites that might contribute to this distribution. Overall, our study does not clearly support either the general-purpose genotype theory or the frozen-niche variation theory for *M. batesii*.

The chemical composition of host plants could also be playing a pivotal role in interactions between *M. batesii* and their host plants as chemistry of host plants frequently exerts a significant influence on herbivory (Howe and Westley 1990; Fritz and Simms 1992; Wittstock and Gershenzon 2002; Nishida 2014; Sánchez-Sánchez and Morquecho-Contreras 2017). Insect herbivores typically display a preference for host plants with low C/N ratios, meaning they contain relatively high nitrogen levels (Strong et al. 1984) including in old field plant communities, arable weeds, and alpine plant communities (Schädler et al. 2003; Karley et al. 2008; Leingärtner et al. 2014). Nitrogen has been shown to be important for growth and development of insect herbivores and therefore has often been used as a proxy for nutritional value of plants (Mattson 1980; White 1984; Wang et al. 2020). Our data suggests a different pattern in *M. batesii*: we found that *M. batesii* in their natural habitats preferred host plants with higher C/N ratios and lower nitrogen levels (Fig. 4). This divergence from the general pattern seen in other species of herbivorous insects suggests that the traits governing herbivore preferences are likely to vary across distinct plant-herbivore interactions (see also Tanentzap et al. 2011). In *M. batesii*, perhaps the preference for higher C/N ratios is due to the synthesis of the alkaloid actinidine. Actinidine is an important component of *M. batesii*’s own chemical defenses; which they synthesize from chemical precursors obtained from eating *Pandanus* and *Benstonea* host-plants (Chow and Lin 1986; Prescott et al. 2009; Vasconcelos et. al., in prep.). The chemical formula for actinidine is C10H13N, suggesting that chemical synthesis of actinidine might impose greater requirements for dietary carbon and hydrogen than for nitrogen. Nitrogen is also a known component of plant defensive compounds, including a defensive compound that serves as the chemical precursor to actinidine (Prescott et al. 2009; Fürstenberg-Hägg et al. 2013). Although *M. batesii* may therefore exhibit a preference for plants containing this defensive chemical, an excess of such compounds in plants could potentially render them toxic to *M. batesii*. Another possibility is that differences in chemical composition observed between chewed and unchewed plants are not a cause but a consequence of chewing by *M. batesii*. Plants often increase production of defensive compounds in response to herbivory (Chen 2008; Wu and Baldwin 2009; Chen and Markham 2021). Ruling out this possibility would require experimental tests. It is important to note that 6 of the 8 uneaten plant samples in the chemical analysis are morphologically very similar, strongly indicating they belong to the same species (*B. lauterbachii*). This suggests that the chemical composition of this species might be the reason it’s avoided in the wild, although we cannot be certain without more samples from different uneaten species.

Ultimately, although we found strong evidence that peppermint stick insects exhibit discernible preferences for a specific host plant chemical composition, and evidence that all-female populations can exploit host plants with tougher leaves, our findings suggest that the host plant phenotypes investigated in this study do not provide a comprehensive explanation for geographical parthenogenesis in *M. batesii*. These results suggest that habitat-related factors, such as host plant morphology and chemistry, may not be the key determinants of sexual or parthenogenetic reproduction in these populations. Instead, this geographical pattern may have resulted primarily from sexual conflict (Burke and Bonduriansky 2018, 2019, 2022; Wilner et al in prep), from stochastic processes such as random dispersal events or male extinction in some mixed-sex populations (see Miller et al. 2024), or from some combination of these factors.

*Megacrania batesii* provides an informative study system for exploring questions regarding geographic parthenogenesis in nature. While certain environmental factors such as host-plant phenotype differentiation cannot explain the patterns observed in the field, there are additional factors to consider in future studies. The presence of parasites, such as bacteria, fungi, biting midges, or parasitic wasps, may offer some explanation for these patterns. Parthenogenetic *M. batesii* populations located in swamp habitats were found to have more ectoparasites on them compared to sexual populations in the same type of environment (Miller et al 2024). It’s possible that certain sites may host a greater number of parasites than others. In environments where parasitic pressure is high, parthenogenetic lineages with limited genetic diversity may struggle to persist over time (i.e. the Red Queen Hypothesis; Bell 1982; Hamilton et al. 1990). Consequently, this could impact the geographic distribution of parthenogenetic lineages across the range. Competition with other insects and predation may also contribute to this. During field observations, we noted orthopterans feeding on the same species of host plants as *M. batesii*, but we did not identify any predators targeting this species. A deeper understanding of the biology and ecology of this system could provide additional clues to the factors that shape sex ratio variation in natural populations. Specifically, further investigation in the differences between sexual and parthenogenetic animals could provide valuable insights into the broader population dynamics of these two coexisting reproductive modes in nature.

## Acknowledgements

We would like to thank Alastair Poore, David Warton, and Ben Walker for advice concerning multivariate statistical analyses. We would also like to thank the Mark Wainwright Analytical Centre Solid State and Elemental Analysis XRF Laboratory for the NCHS analysis used in this study. Lastly, we would like to thank Michele Schiffer for providing access to the JCU Daintree Rainforest Observatory laboratory. This research was funded by the Australian Research Council through Discovery Grant DP200101971 to RB.

Competing Interests Statement:

## The authors declare no competing interests

Statement of Authorship:

Conceptualization: S.M.M., R.B.; funding acquisition: R.B.; data collection: R.B., D.W., J.B., S.M.M.; data analysis and visualization: S.M.M; writing – original draft: S.M.M.; writing – review and editing: R.B., D.W., J.B.

## Data and Code Accessibility

All code is publicly available on Dryad (DOI: 10.5061/dryad.brv15dvhd)

## Appendix

**Figure A1:**
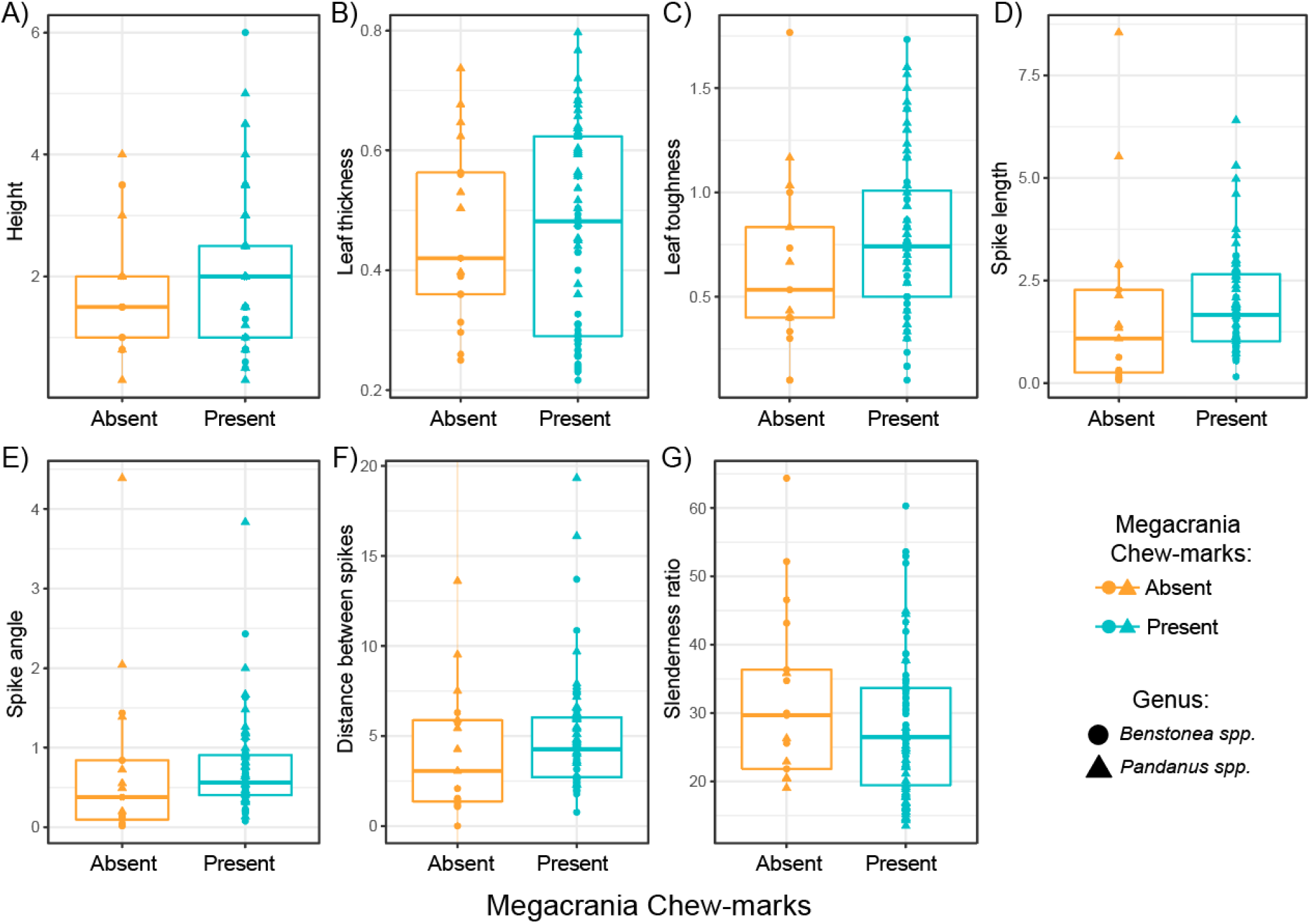
Boxplots showing the effect of host plant morphological traits on *M. batesii herbivory*. Traits include a) height of the plant (m), b) leaf thickness (mm), c) leaf toughness (kg/cm²), d) spike length(mm), e) spike angle (mm distance between end of spike and edge of leaf), f) distance between spikes (mm), and g) slenderness ratio (leaf length/leaf width). The lower and upper hinges of the box correspond to the first and third quartiles (25^th^ and 75^th^ percentiles). The median line is shown. The whiskers extend from the hinges to the largest and smallest value within 1.5 times the inter-quartile range. Outlying points are plotted individually. Colour indicates whether *M. batesii* chew-marks were present or absent.

**Table A1:**
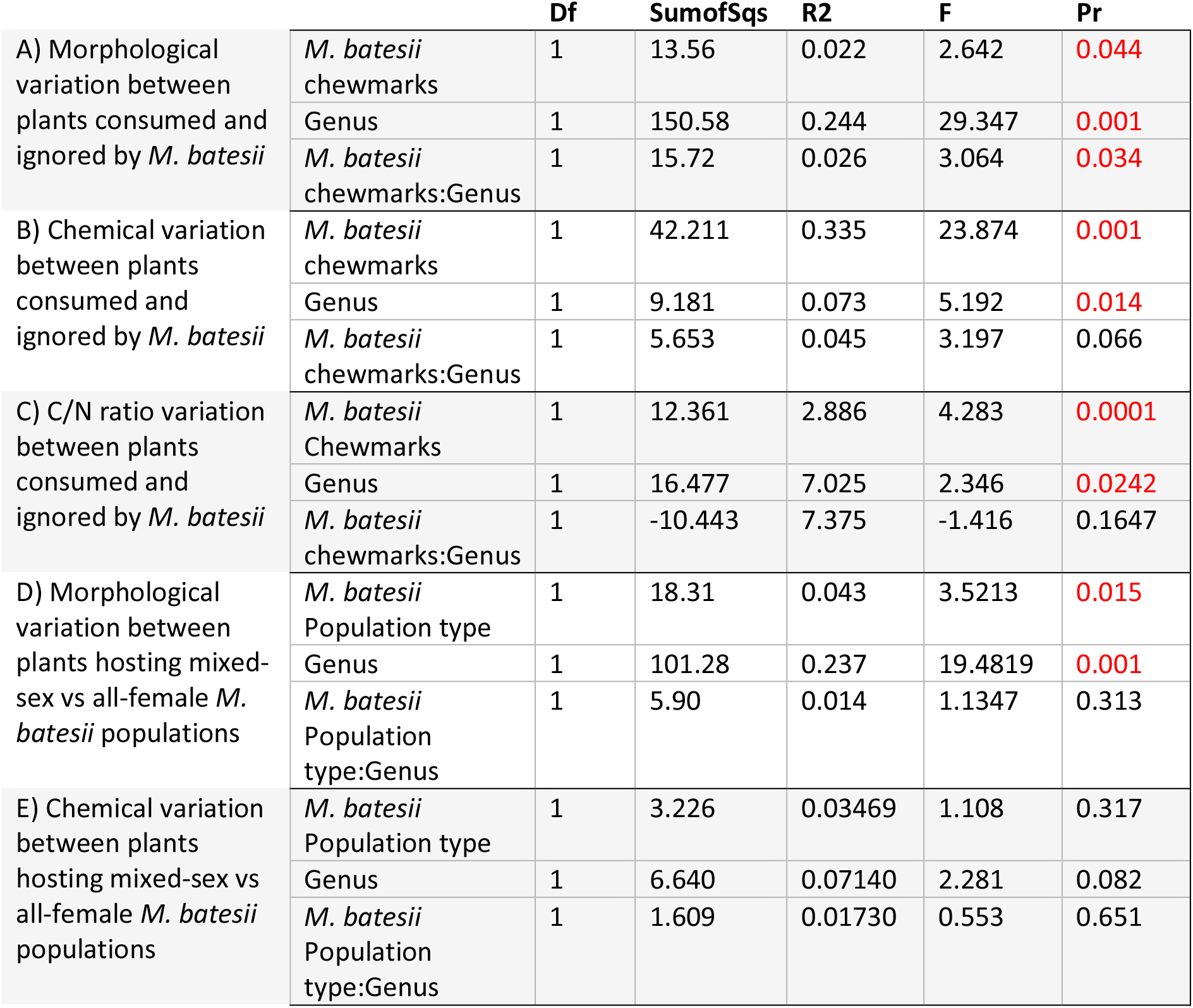
Statistical results for permutational regression tests (A,B,D,E) and linear regression model (C) for A) the morphological variation between plants consumed and ignored by *M. batesii*, B) the chemical variation between plants consumed and ignored by *M. batesii*, C) The C/N ratio variation between plants consumed and ignored by *M. batesii*, D) the morphological variation between plants hosting mixed-sex vs all-female *M. batesii* populations, and E) the chemical variation between plants hosting mixed-sex vs all-female *M. batesii* populations. Df is degrees of freedom, SumofSqs is the sum of squares, R2 is the R-squared value, F is the F-value, and Pr is the p-value associated with the F-value. Significant p-values are colored red.

|  |  | Df | SumofSqs | R2 | F | Pr |
| --- | --- | --- | --- | --- | --- | --- |
| A) Morphological variation between plants consumed and ignored by *M. batesii* | *M. batesii* chewmarks | 1 | 13.56 | 0.022 | 2.642 | 0.044 |
|  | Genus | 1 | 150.58 | 0.244 | 29.347 | 0.001 |
|  | *M. batesii* chewmarks:Genus | 1 | 15.72 | 0.026 | 3.064 | 0.034 |
| B) Chemical variation between plants consumed and ignored by *M. batesii* | *M. batesii* chewmarks | 1 | 42.211 | 0.335 | 23.874 | 0.001 |
|  | Genus | 1 | 9.181 | 0.073 | 5.192 | 0.014 |
|  | *M. batesii* chewmarks:Genus | 1 | 5.653 | 0.045 | 3.197 | 0.066 |
| C) C/N ratio variation between plants consumed and ignored by *M. batesii* | *M. batesii* Chewmarks | 1 | 12.361 | 2.886 | 4.283 | 0.0001 |
|  | Genus | 1 | 16.477 | 7.025 | 2.346 | 0.0242 |
|  | *M. batesii* chewmarks:Genus | 1 | -10.443 | 7.375 | -1.416 | 0.1647 |
| D) Morphological variation between plants hosting mixed-sex vs all-female *M. batesii* populations | *M. batesii* Population type | 1 | 18.31 | 0.043 | 3.5213 | 0.015 |
|  | Genus | 1 | 101.28 | 0.237 | 19.4819 | 0.001 |
|  | *M. batesii* Population type:Genus | 1 | 5.90 | 0.014 | 1.1347 | 0.313 |
| E) Chemical variation between plants hosting mixed-sex vs all-female *M. batesii* populations | *M. batesii* Population type | 1 | 3.226 | 0.03469 | 1.108 | 0.317 |
|  | Genus | 1 | 6.640 | 0.07140 | 2.281 | 0.082 |
|  | *M. batesii* Population type:Genus | 1 | 1.609 | 0.01730 | 0.553 | 0.651 |

**Figure A2:**
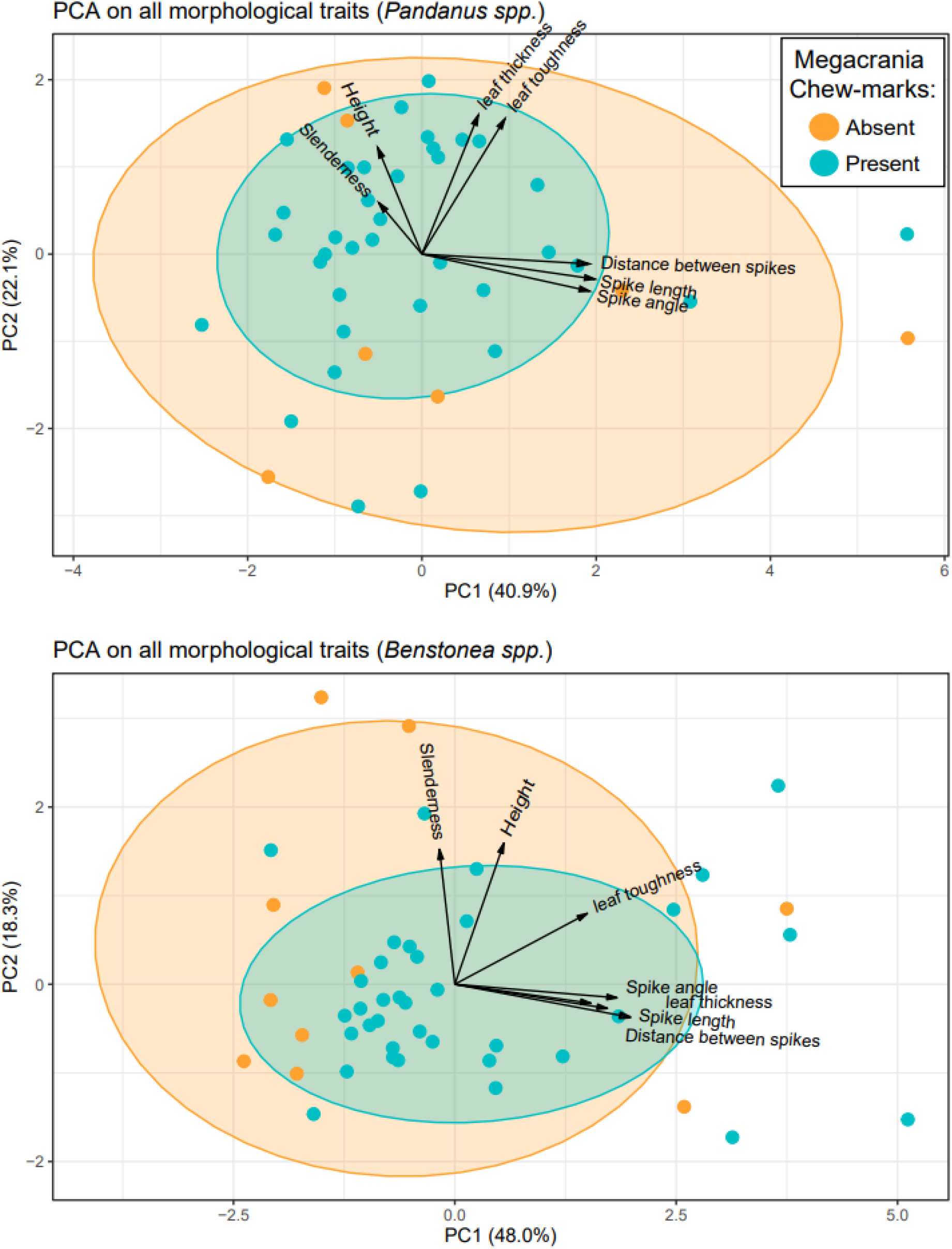
PCA results for host plant morphological traits in a) *Pandanus spp.* and b) *Benstonea spp.* Each point represents an individual host plant sample and color indicates whether *M. batesii* chew-marks were present or absent.

**Table A2:** Coefficients from permutational regression statistical models for the morphological variation between plants consumed and ignored by *M. batesii* in *A) Pandanus spp.* and B) *Benstonea spp.* Columns represent the effect of height of the plant (m), leaf thickness (mm), leaf toughness (kg/cm²), spike angle (mm distance between end of spike and edge of leaf), spike length(mm), distance between spikes (mm), and leaf slenderness ratio (leaf length/leaf width).

|  | Height | Leaf thickness | Leaf toughness | Spike angle | Spike length | Distance between spikes | Slenderness ratio |
| --- | --- | --- | --- | --- | --- | --- | --- |
| A) *Pandanus spp. M. batesii* chewmarks | -0.1885 | -0.0157 | -0.2267 | 0.3224 | 0.2357 | 0.1307 | 0.1243 |
| B) *Benstonea spp. M. batesii* chewmarks | 0.0968 | 0.1416 | -0.1213 | -0.3121 | -0.3434 | -0.2972 | 0.2237 |

**Table A3:** Statistical results for linear mixed-effect models of the effect of population type (Mixed-sex versus all-female) on density of host plants within a 10x10sq m quadrat, with site of the sample as a random effect. For the fixed effects, Estimate is the estimated coefficients, Std.error is the standard errors of the estimated coefficients, df is degrees of freedom, t-value is the t-value associated with the coefficient estimate and Pr is the p-value associated with the t-value. For the random effects, Variance is the variance estimate, and Std.dev is the standard deviation estimates.

| Fixed effects |  |  |  |  |  |
| --- | --- | --- | --- | --- | --- |
|  | Estimate | Std.error | df | t-value | Pr |
| M. batesii Population type | 0.056 | 0.124 | 13.49 | 0.447 | 0.662 |
| Random effects |  |  |  |  |  |
|  | Variance | Std.dev |  |  |  |
| Site | 0.230 | 0.480 |  |  |  |
| Residual | 0.019 | 0.138 |  |  |  |

**Figure A3:**
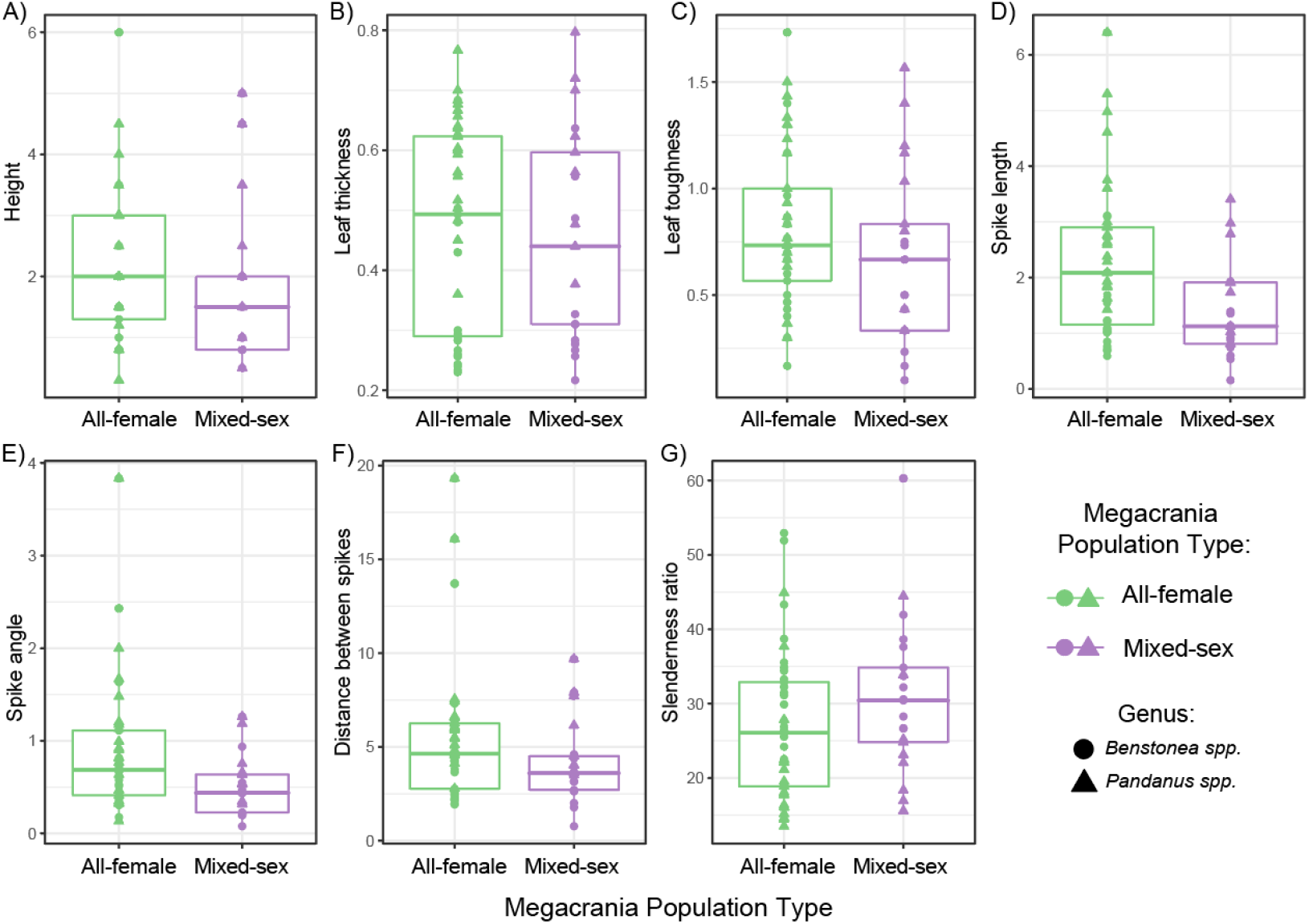
Boxplots showing the relationship between *M. batesii* population type and various host plant morphological traits, including: a) height of the plant (m), b) leaf thickness (mm), c) leaf toughness (kg/cm²), d) spike length(mm), e) spike angle (mm distance between end of spike and edge of leaf), f) distance between spikes (mm), and g) slenderness ratio (leaf length/leaf width). The lower and upper hinges of the box correspond to the first and third quartiles (25^th^ and 75^th^ percentiles). The median line is shown. The whiskers extend from the hinges to the largest and smallest value within 1.5 times the inter-quartile range. Outlying points are plotted individually. Color indicates whether that plant was within an all-female or mixed-sex *M. batesii* population.

**Table A4:**
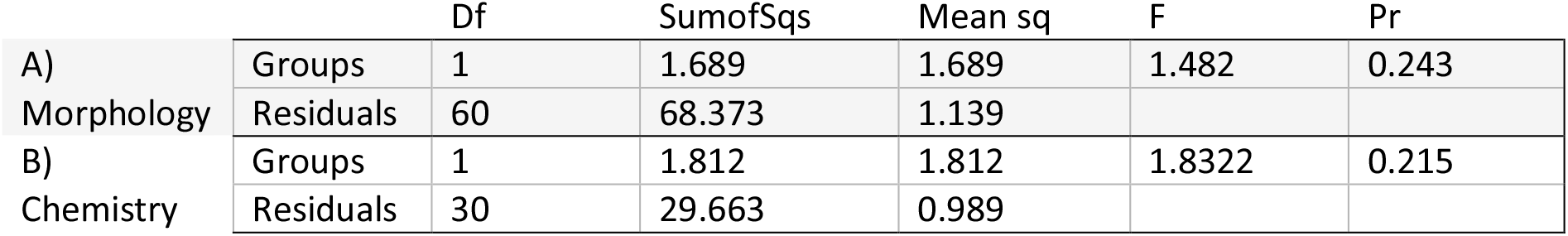
Multivariate homogeneity of groups dispersions test results, testing the differences in group dispersions between habitats hosting all-female *M. batesii* populations and mixed-sex *M. batesii* populations for a) morphological phenotypic traits and b) chemical phenotypic traits. Df is degrees of freedom, SumofSqs is the sum of squares, Mean sq is the squared mean, F is the F-value, and Pr is the p-value associated with the F-value.

**Table A5:** Statistical results for linear mixed-effect models of the effect of population type (Mixed-sex versus all-female) on group dispersions, with site of the sample as a random effect. For the fixed effects, Estimate is the estimated coefficients, Std.error is the standard errors of the estimated coefficients, df is degrees of freedom, t-value is the t-value associated with the coefficient estimate and Pr is the p-value associated with the t-value. For the random effects, Variance is the variance estimate, and Std.dev is the standard deviation estimates.

| Fixed effects |  |  |  |  |  |
| --- | --- | --- | --- | --- | --- |
|  | Estimate | Std.error | df | t-value | Pr |
| *M. batesii* Population type | -0.1545 | 0.300 | 11.2 | -0.515 | 0.617 |
| Random effects |  |  |  |  |  |
|  | Variance | Std.dev |  |  |  |
| Site | 0.200 | 0.447 |  |  |  |
| Residual | 0.352 | 0.593 |  |  |  |

B) Chemical
| Fixed effects |  |  |  |  |  |
| --- | --- | --- | --- | --- | --- |
|  | Estimate | Std.error | df | t-value | Pr |
| *M. batesii* Population type | 0.158 | 0.205 | 6.11 | 0.771 | 0.469 |
| Random effects |  |  |  |  |  |
|  | Variance | Std.dev |  |  |  |
| Site | 0.041 | 0.204 |  |  |  |
| Residual | 0.153 | 0.392 |  |  |  |

